# Harmonized nonhuman cancer database

**DOI:** 10.1101/2025.10.19.683290

**Authors:** Eslam Abousamra, Benjamin Jagt, Esther Osafo, Nash D. Rochman

## Abstract

All multicellular metazoa, including both vertebrates and invertebrates, are susceptible to cancer. Cancer prevalence among vertebrate species ranges from less than 1% to more than 50% and is dependent on tissue type. Species generally more or less susceptible to cancer overall may exhibit a low or high rate of a specific tumor. Understanding the genetic or lifestyle factors which impact the relative rate of tumorigenesis in different tissues across species would support new approaches to personalized and preventative therapy. Research in this area is challenging due to heterogeneous reporting criteria including inconsistent and species-specific tumor metadata. Here we present a harmonized dataset including tumors from 41,539 individuals across 825 species. Each tumor is assigned the NCBI taxonomy ID for the species where it was found; a tissue-type ID specifying body site if applicable; and a cell type ID. Using this dataset, we demonstrate the relative incidence of breast, colon, and lung cancers among nonhuman species is typically low while soft tissue relative incidence is high. These results suggest the existence of unrecognized genetic machinery involved in tumor suppression or oncogenesis which may be key drivers of non-human cancer and likely play a role in human disease progression, including metastasis.

## Introduction

The lifetime risk for an invasive cancer diagnosis in the United States is 41.6% for men and 39.6% for women. Despite enormous research investment, cancer also remains the second leading cause of death^1^. Reducing overall cancer mortality is challenging due in large part to the varied genetic etiology of tumors originating from different tissue types^2^. The factors shaping tumor evolution in response to treatment are also dependent on tissue type^3^ as well as germline genetic variation^4, 5^. Tumor mutation databases, notably The Cancer Genome Atlas (TCGA)^6^ and The Catalogue of Somatic Mutations in Cancer (COSMIC)^7^, have revealed conserved evolutionary signatures suitable for patient subtyping and personalized therapy^8, 9^. These resources remain fundamentally limited, however, by the relatively homogeneous genetic background of the, human, patient population. Extending these efforts to other species would allow for more robust statistical analysis of the relationships between germline genetic variation and tumor evolution potentially revealing protective or deleterious genetic features which are pervasive in other species but conditionally dependent in humans.

All multicellular metazoa, including both vertebrates and invertebrates, are susceptible to cancer^10^. Cancer has been identified across both extant species and within the fossil record^11-13^. A major focus of evolutionary and comparative oncology has been to reveal the factors underpinning the variance in cancer prevalence across species. Cancer prevalence among vertebrate species ranges from less than 1% to more than 50%^14^ and depends on both lifestyle and genetic factors which remain relatively poorly understood. Classically, adult body size had been assumed to be associated with higher rates of cancer as larger body size corresponds to a greater total number of cell divisions during development and tissue homeostasis. Early efforts to quantify this trend arrived at the contrary conclusion that body size is independent or negatively correlated with cancer prevalence, often referred to as Peto’s Paradox^15^. This long standing result may be due in part to observational bias towards a low diversity of charismatic large species and familiar or lab-friendly small species^15-19^. More recently, the identification of diverse small species with extremely low rates of cancer including notably several species of bat and the naked mole-rat^20-22^ motivated a reassessment of the relationship between body size and cancer prevalence. The most comprehensive analysis of vertebrate cancer prevalence available at the time of writing has demonstrated body size to be positively correlated with cancer prevalence overall^14^, although the trend within mammals may be less clear^17^.

These results are in agreement with the finding that, among individuals of the same species, cancer risk scales with body size^15^. Relatedly, litter and clutch size in mammals and birds respectively^17, 23, 24^ is positively correlated, and gestation time^14^ is negatively correlated with cancer prevalence. In addition to observational bias, another possible explanation for why resolving the relationship between body size and cancer prevalence continues to be so challenging is that each stem cell division may confer substantially greater risk of tumorigenesis than any somatic cell division^25^. The total number of stem cell divisions during an individual’s life may also be correlated with litter/clutch size. Consequently, tissues with low rates of stem cell division (e.g. adipose tissue or skeletal muscle) which comprise the majority of the body mass of many species may not confer substantially increased cancer risk.

Here we consider a tissue-specific approach to evaluate the relative incidence of different cancer subtypes arising in different body sites. We analyze a dataset including a large number of case reports for which the number of individuals under observation is not known (see Fig. 1A for an illustration). This prohibits the measurement of prevalence but expands the diversity of species studied. Through this approach, we are able to provide harmonized NCBI taxonomy IDs for the species in which the tumor was found; tissue-type IDs specifying body site where appropriate; and cell type IDs for tumors observed within approximately 1000 species, including major taxonomic groups for which controlled zoological data is unavailable. Supported with this data, we demonstrate the relative incidence of breast, colon, and lung cancers among nonhuman species is typically low while soft tissue relative incidence is high. These species-specific trends may be attributed to the existence of unrecognized genetic machinery involved in tumor suppression or oncogenesis which may also play a role in human disease progression.

**Figure 1.**
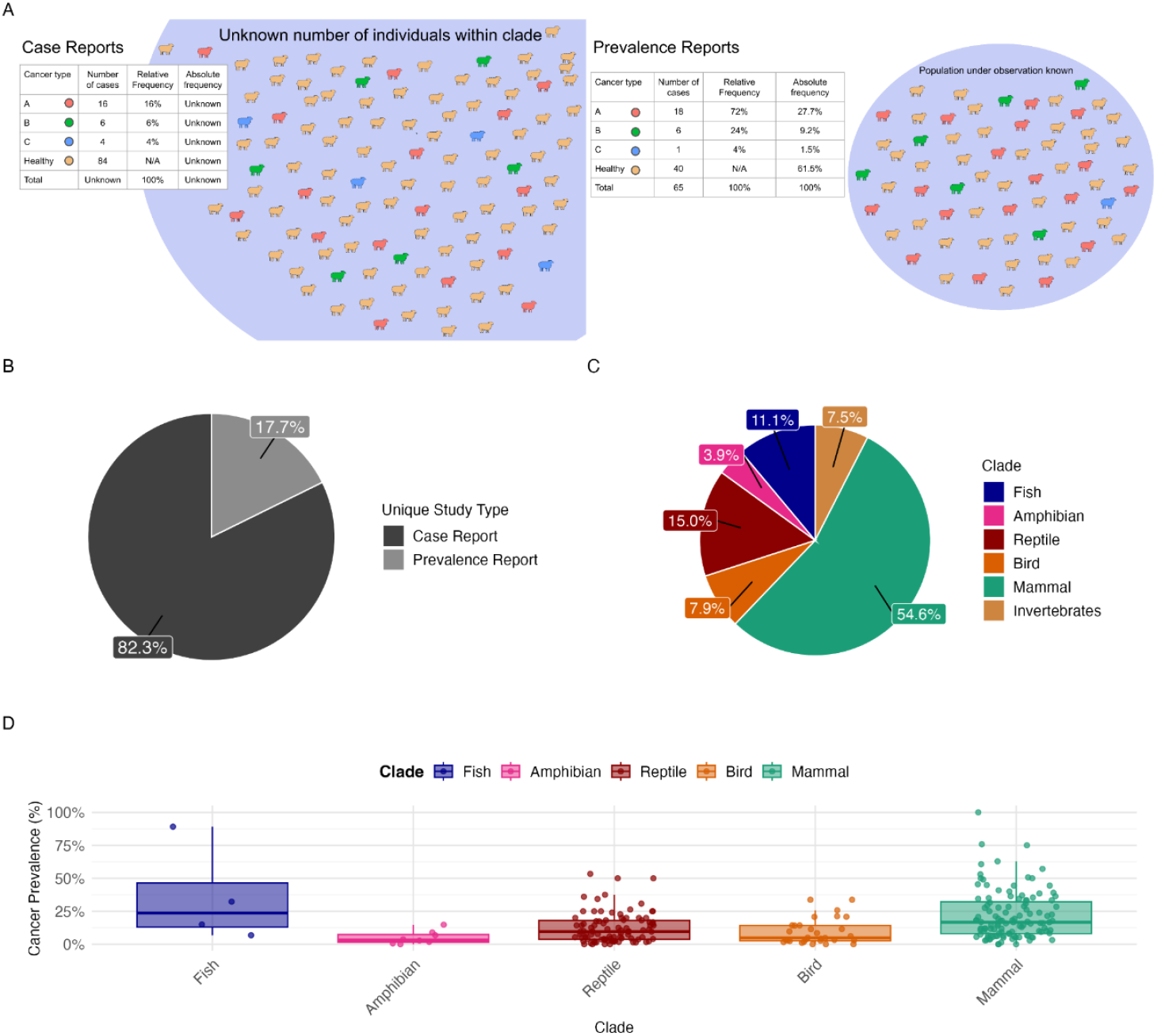
Despite focus on human disease and economics, available data on cancer in diverse species is extensive. **A:** We divide the literature into 2 categories: Prevalence Reports, conducted over populations for which the total number of individuals is known, and Case Reports for which the population under observation is poorly characterized. Case Reports support the estimation of relative frequencies of cancer subtypes but not prevalence. **B:** Categorization of nonhuman cancer research literature. **C:** Distribution of species included in Prevalence Reports. **D:** Cancer prevalence across major taxa where each point is a species.

## Results

Our database is available for download here: https://zenodo.org/records/17391660. We sought to establish a harmonized database of peer-reviewed reports of nonhuman cancer across both invertebrates and vertebrates. To assemble this database, we conducted a network-assisted literature search utilizing ResearchRabbit (researchrabbit.ai/). For most taxonomic groups, this search strategy was exhaustive such that the network was extended until no additional articles could be identified with matching keywords or connections in the citation graph. For select, well-studied, taxonomic groups, principally lab animals, this search strategy was not exhaustive. See Methods for details. Expectedly, data availability was greater for species with greater perceived relevance for human disease or economics.

### Inclusion of Case Reports expands species diversity

Prior work can be coarsely divided into six categories. 1) Transmissible cancers including those affecting canines^26, 27^ which are both companion animals and viable models for human disease, Tasmanian devils^26, 28-32^, mollusks^33-35^, and hydra^36, 37^ (vertical transmission). 2) Model species including primarily primates^38, 39^, mice^40^, rats^41-44^, hamsters^45, 46^, ferrets^47^, and zebrafish^48^. 3) Companion animals including both transmissible and nontransmissible cancers in dogs^49, 50^, cats^51^, ferrets, rabbits, and guinea pigs^52^. 4) Livestock including primarily ruminants, swine^53^, and chickens^54^ (in particular, hyper virus related papilloma^55^). 5) Zoo populations, many of which are well-controlled where the total numbers of individuals under observation are known^10, 14, 56^. 6) Wild populations, which are almost exclusively uncontrolled case reports and tend to focus on unusual subpopulations including notably the St. Lawrence beluga whales^57, 58^ and groups of Bighorn Sheep^59^ which are likely impacted by localized environmental carcinogens.

Despite this historical focus on select species and cancer subtypes, the overall data availability of nonhuman cancer is robust. Our dataset includes 48 controlled zoological studies where the number of individuals under observation is known, classified as Prevalence Reports, and 223 case Reports where it is not known (Fig. 1B). Many Prevalence Reports focus only on mammalian cancer (Fig. 1C); however, prevalence across all major taxa is high enough to support the statistical analysis of body site variation (Fig. 1D). Our dataset covers 41,539 individuals across 825 species. Note that subpopulations known to be impacted by localized environmental carcinogens are included in the dataset but excluded from all analysis presented in the manuscript. Please see Supplementary Data 1.

The inclusion of Case Reports substantially increases both the number of publications (Fig. 2A) and tumors (Fig. 2B) contributing to the data set. Notably, several major clades are overwhelmingly represented by Case Reports, including most invertebrates (Protostomia), 60% of all bird species (Passeriformes), iguanas and turtles (Iguania and Testudinata), sharks and rays (Chondrichthyes), and whales (Whippomorpha). Case Report data remains limited, however, because it is difficult or impossible to measure cancer prevalence utilizing only Case Reports. Individuals without cancer are likely to go unreported and, overall, the prevalence of cancer is likely to be overestimated.

**Figure 2.**
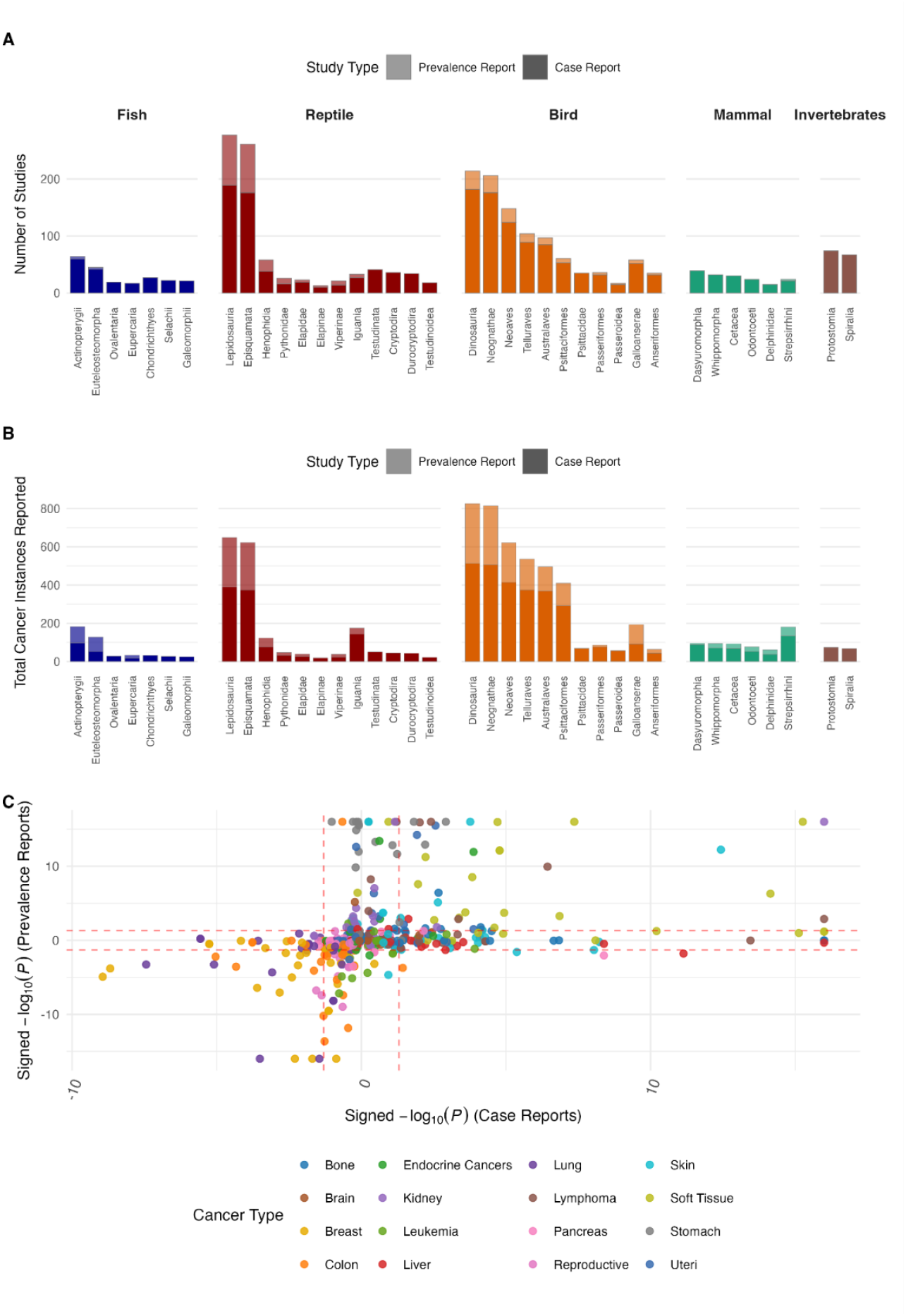
Inclusion of Case Reports improves species coverage. **A:** Study counts across major taxa where analysis is primarily facilitated by the inclusion of Case Reports. See methods for selection of taxa. **B:** Individual tumor counts across taxa in A. **C:** Signed likelihood that each tumor category is more (+) or less (-) common in each clade relative to the human reference. Likelihoods computed independently utilizing only Case Reports (x-axis) or Prevalence Reports (y-axis). Breast shown only for nonhuman mammals. Dashed lines indicate P=0.05.

In principle, Case Report data can be used to compute the relative incidence of different tumor types. We classify tumors into 16 tissue-type categories: Bone, Brain, Breast, Colon, Endocrine Cancers, Kidney, Leukemia, Liver, Lung, Lymphoma, Pancreas, Reproductive, Skin, Soft Tissue, Stomach, and Uteri. Additionally, we assign each tumor a cell type category: carcinomas derived from epithelial cells; sarcomas derived from mesenchymal cells in bone or connective tissues; lymphohematopoietic cancers derived from mesenchymal cells in blood or lymphatic tissues; and benign tumors (assigned irrespective of their apparent cell type of origin). Benign tumors are assigned a tissue-type category but excluded from the analysis. Despite heterogeneous reporting criteria including inconsistent and species-specific tumor metadata, approximately 90% of all tumors can be assigned harmonized tissue- and cell-type labels. This roughly 90% characterization rate is consistent across major taxonomic groups (Fig. S1). Smaller studies and Case Reports are no less well characterized than large Prevalence Reports (Fig. S2).

It is plausible, however, that Case Report data remains subject to more significant observation or reporting bias than Prevalence Report data. For example, it may be anticipated that surface tumors or tumors most commonly observed among humans would be overrepresented in Case Reports of wild populations relative to necropsy data from the same species in a zoological park. To assess this, we computed the relative incidence of each of the 16 tissue-type categories among the 32 taxonomic groups representing the shallowest taxonomic groups containing at least 10 species contributing at least one tumor in the dataset (selected moving from tip to root within the species tree). For every group, *n*, and each category, *i*, we compute the probability of observing *n*_*i*_ or more/fewer tumors of category *i* in an ensemble of *Σ*_*i*_*n*_*i*_ randomly selected human tumors if that category was over/under represented in that clade relative to the human reference, respectively (see Methods for details).

Figure 2C displays the signed likelihood that each tumor category is more or less common in each group relative to the human reference where the likelihoods were computed twice independently utilizing only Case Reports or Prevalence Reports. Only 4 of the 512 (32 groups, 16 categories) display significant but opposing deviations from the human reference, demonstrating that the Case Report data is consistent with data produced by more tightly regulated Prevalence Reports. In direct opposition to the anticipated humanizing observation or reporting bias, three out of four of the most common sites in terminal human cancer, Breast (only mammals shown), Colon, and Lung, are significantly underrepresented across most groups. This finding is consistent with human-specific physiological and environmental factors (see Discussion). Conversely, soft tissue cancer is significantly overrepresented among nonhuman species. Soft tissue cancers have a heterogenous tissue and body site presentation, but nearly all soft tissue tumors are sarcomas, and indeed among human tumors, soft tissue cancer classification is often synonymous with sarcoma.

### Sarcomas are overrepresented among nonhuman tumors

Most human cancers fall into one of three primary groups: carcinomas derived from epithelial cells; sarcomas derived from mesenchymal cells in bone or connective tissues; and lymphohematopoietic cancers derived from mesenchymal cells in blood or lymphatic tissues. Most human cancers are carcinomas, followed by lymphohematopoietic cancers, while sarcomas tend to be rare. Sarcomas represent less than 1.0% of adult human cancers but a much larger fraction of cancers observed in children, 20%^62^. Mean sarcoma incidence is also approximately 20% across groups of fish and reptiles with more significant variability among reptiles. While still 10-fold higher than adult humans, sarcoma incidence is lower among birds, approximately, 10% for most groups, but very high among the parrots represented in this dataset (Psittacidae, >50%). Sarcoma incidence is most variable among mammals ranging from over 40% among the family containing horses and rhinos (Perissodactyla) to no cases at all among dolphins (Delphinidae). This variance does not appear to be explainable by commonly computed life history characteristics (e.g. body size, lifespan, or litter size), see Discussion.

As introduced in the previous section, not all tumors have metadata consistent with this three-class cancer categorization including benign tumors independent of the cell type of origin and tumors stemming from other progenitor cells (in particular, many tumors identified within reproductive organs). Fig. 3B displays the relative proportion of the three primary categories. Like sarcoma, major groups are observed to have substantially elevated lymphohematopoietic cancer incidence in comparison to humans.

**Figure 3.**
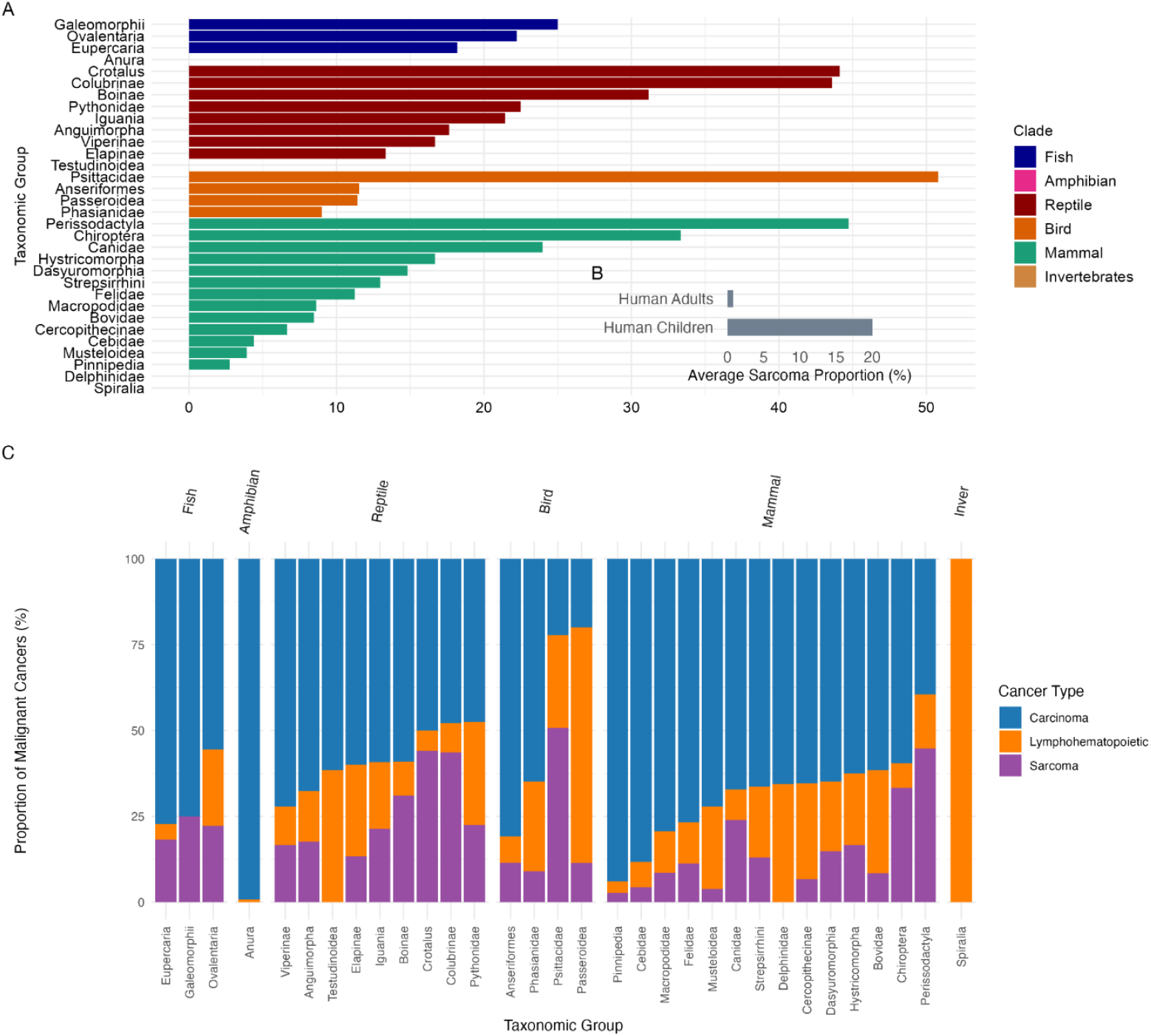
Relative incidence of cancers categorized by cell type of origin. Proportion of tumors which are sarcomas. **B:** Relative proportion of carcinomas, sarcomas, and lymphohematopoietic tumors, excluding other tumors categories (e.g. benign).

## Discussion

Here we present a curated dataset merging 271 studies of nonhuman cancer among 41,539 individuals across 825 species (see Supplementary Table 1). Mammals constitute the largest fraction of the species, followed by Reptiles, Fish, Birds, Invertebrates, and Amphibians. This dataset includes 48 controlled zoological studies where the number of individuals under observation is known, Prevalence Reports, and 223 Case Reports where it is not known. No systematic bias, humanizing or otherwise, among Case Reports was identified. To the contrary, it is also possible that the individuals studied in Prevalence Reports, which primarily reside within zoological parks, may demonstrate different cancer phenomenology than their wild counterparts due to the specific environmental conditions experienced in captivity. Species with a successful history of animal husbandry^63^ are also enriched within zoo populations. Overall, we believe our assembly and metadata harmonization of these studies has produced a robust and reusable dataset which we hope will motivate reuse and integration into future efforts.

Our initial analysis of this dataset produced 4 principal findings: breast, colon, and lung cancers are significantly underrepresented among nonhuman tumors relative to humans while sarcomas are significantly overrepresented. These underrepresented tumor types have straightforward potential explanations. Permanent, adipose breasts are a uniquely human trait, with complex hormonal regulation^64^ that likely influences tumorigenesis. The majority of lung cancers are attributed to uniquely human environmental exposures, principally smoking^65^. Medical support extending the human lifespan increases the prevalence of all age-associated disease, including cancer. The relative incidence of the most common cancer types, which are driven by biological processes that persist across the adult lifespan and are not primarily limited to developmental or reproductive events, are the most likely to be influenced by these human-specific modifiers of the age distribution. Colon epithelial cells continuously turnover at a high rate^66^ and are subject to intense environmental stress. Additionally, colon cancer may be driven in part by human-specific environmental exposures^65^.

The relatively higher rate of sarcoma across most nonhuman groups is not as clearly and immediately explainable. Previous work has suggested a carnivorous diet is associated with increased sarcoma risk^10^ including a sarcoma proportion of ∼12-15% in canines^67, 68.^. We also observe a relatively high rate of sarcoma among canines (Canidae, ∼25%) as well as cats (Felidae, ∼12%); however, we find low relative rates of sarcoma, <5%, among the families containing weasels, seals, and dolphins (Musteloidea, Pinnipedia, and Delphinidae).

As with colon cancer, prolonged post-developmental or post-reproductive lifespan is another potential explanation for the relatively low rates of sarcoma among human populations. Sarcomas represent less than 1.0% of adult human cancers but a much larger fraction of cancers observed in children, 20%^62^. This does not immediately explain the variance among nonhuman species, however. Bats, the longest-lived mammals when adjusted for body size^69^, and parrots, which also exhibit a long body-size adjusted lifespan^70^, display among the highest relative incidence of sarcoma (Chiroptera, ∼35% and Psittacidae, >50%, respectively). One potential explanation for this variance is species-specific body composition diversity, significantly altering the proportion of different progenitor stem cell numbers, which may have an outsized impact on the risk of tumorigenesis in comparison to somatic cell divisions^25^.

Another possibility is species-specific variation among cancer driver genes. Our previous work demonstrated that, on the whole, human cancer driver mutations are strongly deleterious across all cellular life wherever orthologous sites can be identified^61^. This includes a large number of driver mutations predicted to be pathogenic in multicellular Plants and Fungi which are not subject to cancer as well as a smaller ensemble of drivers which are likely pathogenic even among, single-celled, prokaryotes. Consequently, it may be expected that many, if not the majority, of nonhuman cancers are driven in part by established human drivers. Notably, variation in the copy number of the human driver gene, tumor suppressor TP53 has been implicated in modulating cancer prevalence among Mammals^71^ and species with low rates of cancer, including elephants^72^ and the naked mole rat^73^ tend to have many copies of TP53.

However, we also uncovered a significant ensemble of human cancer driver mutations which are the consensus residue among major taxonomic groups, even including some mammals. Most, but not all, instances appear to be subject to compensatory epistasis such that the driver co-occurs with another substitution relative to the human reference, in some cases another driver mutation. With or without a compensatory residue, the presence of these driver mutations likely alters the genomic etiology of tumorigenesis and tumor evolution in these species. These variants, or indeed mutations in genes that are currently not identified as human oncogenes or tumor suppressors, may act as conditional drivers^74^ during metastasis in the context of previously acquired mutations in the primary tumor. Uncovering such individual variation is essential for designing personalized therapy.

Recent work dramatically expanding the diversity of available vertebrate reference genomes is likely to support the identification of new cancer-associated genes^75^; however, without investment in nonhuman tumor sequencing, differentiating between protective, pathogenic, and neutral genomic features will remain extremely challenging. We hope this established finding, that a significant ensemble of human cancer driver mutations are the consensus residue among major taxonomic groups, coupled with the results of the present study demonstrating that the significant diversity in cancer phenotypes across vertebrates is not immediately explainable by variation in well-studied life-history traits, will motivate serious investment in nonhuman tumor sequencing.

## Data Availability

Our database is available for download here: https://zenodo.org/records/17391660.

## Methods

### Data Collection

We conducted a network-assisted literature review using Research Rabbit^76^. Initial nodes in the network were seeded with articles matching specified keywords. Moving from root to tip within the metazoan species tree, the citation graph was explored until apparent saturation for most taxonomic groups (no new articles citing, referencing, or with shared authorship were found for any node in the network reporting tumors within that taxonomic group). An ad-hoc literature review for taxonomic groups which yielded few cancer reports was independently conducted by a second reviewer and in every case did not yield any additional relevant articles. Our network search was ended prior to saturation for a minority of extensively studied taxonomic groups, principally lab animals. Studies were included in our dataset only if the tumors reported were apparently naturally occurring. Subpopulations known to be impacted by localized environmental carcinogens are included in the dataset but excluded from all analysis presented in the manuscript. All data are presented in Supplementary Data 1.

### Taxonomic Grouping

Heterogeneity in sampling prevented reporting statistics for a conserved taxonomic rank (e.g. species) with some taxonomic groups subject to a dramatically higher sampling density than others. Consequently, we constructed taxonomic groups of variable rank, selected to be as narrow as possible while still containing a suitable number of distinct species with at least one tumor observed. Moving from leaf to root, following the NCBI taxonomy tree, the total number of descendent species contributing at least one tumor was computed. The shallowest taxonomic group containing tumors from 10 distinct species was selected resulting in 32 such nonoverlapping taxonomic groups. Species for which the shallowest ancestral group containing at least 10 species also contains a parallel descendent group with at least 10 species are not included in this grouping. This grouping is applied in Fig. 2C and Fig. 3.

Fig. 2A-B includes all taxonomic groups meeting the following criteria. Each group must contain 10 species with at least 1 tumor. Each group must be in the top 50 such groups ordered by the fraction of species within that group that are only represented by Case Reports or the fraction of tumors within that group that come from Case Reports. When more than one NCBI taxonomy ID was consistent with the species selected, the shallowest NCBI taxonomy ID was retained (e.g. if all representatives of the group “green and blue birds” are also representative of the group “green birds” the group would be assigned the NCBI taxonomy ID for “green birds”. Lastly the basal groups Sauropsida, Archosauria, and Archelosauria were removed, all of which are represented by highly skewed distributions of descendent species within the dataset and not reflecting their names. This procedure yielded 38 groups.

Across all figures, taxonomic groups are identified by common names of major groupings: fish, reptiles, amphibians, mammals, birds, and invertebrates. These labels were assigned to all descendants of the following clades. Note that while Protostomia does not encompass all species commonly called “invertebrates” it does encompass all such representatives in this dataset.

Fish: Actinopterygii, Dipnomorpha, Coelacanthimorpha, Dipnomorpha, Chondrichthyes Reptiles: Testudinata, Crocodylia, Lepidosauria

Amphibians: Amphibia Mammals: Mammalia Birds: Dinosauria Invertebrates: Protostomia

### Tumor Classification

Each tumor was classified in 2 ways: tissue-type / body site harmonized to match to 16 cancer types, including the most affected organ systems among human cancer mortalities^77^, and cell type of origin.

Tissue-type / body site was assigned as follows. First, Hematopoietic/lymphoid malignancies were subdivided into Leukemia or Lymphoma categories using expanded keyword rules (e.g., “myeloid” to Leukemia; “hodgkin” to Lymphoma). Next unmatched terms were mapped to 14 organ specific categories (Breast, Colon, Kidney, Liver, Lung, Pancreas, Stomach, Uteri, Brain, Bone, Skin, Soft Tissue, Reproductive, Endocrine Cancers) through anatomical keyword patterns, (e.g., “hepat” to Liver; “pulmon” to Lung), with clade-specific adjustments, and finally designating unclassifiable cases as “Other.” We then approximated human cancer distributions using occurrence data with human cancer probabilities assigned to each cancer category.^89-90^

Cell type of origin classification was performed using comprehensive keyword dictionaries to achieve assignments to one of 4 categories: carcinoma, sarcoma, lymphohematopoietic, benign, or other. First, Hematopoietic/lymphoid malignancies were consolidated into a single lymphohematopoietic category via exact keyword matching (e.g., “lymphoma,” “leukemia,” “plasmacytoma”), prioritizing cellular lineage over anatomic site. Next, remaining terms were systematically mapped to predefined dictionaries of histopathological terms—such as “adenocarcinoma” (carcinoma), “leiomyosarcoma” (sarcoma), or “adenoma” (benign)—using regular expressions to account for morphological variants and plurals. Ambiguous cases - those matching multiple categories or partially overlapping patterns - were resolved through a sequential rule hierarchy prioritizing hematopoietic lineage over mesenchymal, and mesenchymal over epithelial origin. Remaining unmatched or morphologically indeterminate terms (e.g., “blastoma,” “undifferentiated neoplasm,” or unspecified “tumor”) were assigned to other.

### Evaluating significance in deviations from the human reference

In order to evaluate statistically significant variation between the relative incidence of different tumor types across taxonomic groups, we first evaluated whether each tumor type was significantly over or underrepresented relative to the established human reference cancer distribution for occurrence^72,89-90^. For every clade, *n*, and each category, *i*, we computed the probability of observing *n*_*i*_ or more/fewer tumors of category *i* in an ensemble of *Σ*_*i*_*n*_*i*_ randomly selected human tumors if that category was over/underrepresented in that clade relative to the human reference.

## Acknowledgements

We thank Yuri Wolf and members of the Evolutionary Health Group for the valuable discussions. NDR is supported through the Intramural Research Program of the National Library of Medicine, National Institutes of Health and by intramural support from the City University of New York Graduate School of Public Health and Health Policy. The contributions of the NIH authors are considered Works of the United States Government. The findings and conclusions presented in this paper are those of the authors and do not necessarily reflect the views of the NIH or the U.S. Department of Health and Human Services.

## References

1. Siegel RL, Giaquinto AN, Jemal A. Cancer statistics, 2024. CA: A Cancer Journal for Clinicians. 2024; 74:12–49.

2. Haigis KM, Cichowski K, Elledge SJ. Tissue-specificity in cancer: The rule, not the exception. Science. 2019; 363:1150–1.

3. McGranahan N, Swanton C. Clonal Heterogeneity and Tumor Evolution: Past, Present, and the Future. Cell. 2017; 168:613–28.

4. Ramroop JR, Gerber MM, Toland AE. Germline Variants Impact Somatic Events during Tumorigenesis. Trends Genet. 2019; 35:515–26.

5. Chatrath A, Ratan A, Dutta A. Germline Variants That Affect Tumor Progression. Trends in Genetics. 2021; 37:433–43.

6. Weinstein JN, Collisson EA, Mills GB, Shaw KRM, Ozenberger BA, Ellrott K, et al. The Cancer Genome Atlas Pan-Cancer analysis project. Nature Genetics. 2013; 45:1113–20.

7. Tate JG, Bamford S, Jubb HC, Sondka Z, Beare DM, Bindal N, et al. COSMIC: the Catalogue Of Somatic Mutations In Cancer. Nucleic Acids Research. 2019; 47:D941–D7.

8. Uzilov AV, Ding W, Fink MY, Antipin Y, Brohl AS, Davis C, et al. Development and clinical application of an integrative genomic approach to personalized cancer therapy. Genome Medicine. 2016; 8.

9. Clayton EA, Pujol TA, McDonald JF, Qiu P. Leveraging TCGA gene expression data to build predictive models for cancer drug response. BMC Bioinformatics. 2020; 21.

10. Vincze O, Colchero F, Lemaître J-F, Conde DA, Pavard S, Bieuville M, et al. Cancer risk across mammals. Nature. 2022; 601:263–7.

11. Odes EJ, Randolph-Quinney PS, Steyn M, Throckmorton Z, Smilg JS, Zipfel B, et al. Earliest hominin cancer: 1.7-million-year-old osteosarcoma from Swartkrans Cave, South Africa. South African Journal of Science. 2016; 112:5.

12. Rothschild BM, Tanke DH, Helbling M, Martin LD. Epidemiologic study of tumors in dinosaurs. Naturwissenschaften. 2003; 90:495–500.

13. Haridy Y, Witzmann F, Asbach P, Schoch RR, Frobisch N, Rothschild BM. Triassic Cancer-Osteosarcoma in a 240-Million-Year-Old Stem-Turtle. JAMA Oncol. 2019; 5:425–6.

14. Compton Z, Harris V, Mellon W, Rupp S, Mallo D, Kapsetaki SE, et al. Cancer Prevalence Across Vertebrates. 2023.

15. Caulin AF, Maley CC. Peto’s Paradox: evolution’s prescription for cancer prevention. Trends in Ecology & Evolution. 2011; 26:175–82.

16. Gaughran SJ, Pless E, Stearns SC. How elephants beat cancer. Elife. 2016; 5.

17. Boddy AM, Abegglen LM, Pessier AP, Aktipis A, Schiffman JD, Maley CC, et al. Lifetime cancer prevalence and life history traits in mammals. Evol Med Public Health. 2020; 2020:187–95.

18. Lair S, Barker IK, Mehren KG, Williams ES. Epidemiology of neoplasia in captive black-footed ferrets (Mustela nigripes), 1986-1996. J Zoo Wildl Med. 2002; 33:204–13.

19. Newman SJ, Smith SA. Marine mammal neoplasia: a review. Vet Pathol. 2006; 43:865–80.

20. Taylor KR, Milone NA, Rodriguez CE. Four Cases of Spontaneous Neoplasia in the Naked Mole-Rat (Heterocephalus glaber), A Putative Cancer-Resistant Species. The Journals of Gerontology Series A: Biological Sciences and Medical Sciences. 2017; 72:38–43.

21. Delaney MA, Ward JM, Walsh TF, Chinnadurai SK, Kerns K, Kinsel MJ, et al. Initial Case Reports of Cancer in Naked Mole-rats (Heterocephalus glaber). Vet Pathol. 2016; 53:691–6.

22. Hua R, Ma Y-S, Yang L, Hao J-J, Hua Q-Y, Shi L-Y, et al. Experimental evidence for cancer resistance in a bat species. Nature Communications. 2024; 15.

23. Kapsetaki SE, Compton Z, Dolan J, Harris VK, Rupp SM, Duke EG, et al. Life history and cancer in birds: clutch size predicts cancer. 2023.

24. Dujon AM, Vincze O, Lemaitre J-F, Alix-Panabières C, Pujol P, Giraudeau M, et al. The effect of placentation type, litter size, lactation and gestation length on cancer risk in mammals. Proceedings of the Royal Society B: Biological Sciences. 2023; 290.

25. Tomasetti C, Vogelstein B. Cancer etiology. Variation in cancer risk among tissues can be explained by the number of stem cell divisions. Science. 2015; 347:78–81.

26. Siddle HV, Kaufman J. Immunology of naturally transmissible tumours. Immunology. 2015; 144:11–20.

27. Murchison EP, Wedge DC, Alexandrov LB, Fu B, Martincorena I, Ning Z, et al. Transmissible [corrected] dog cancer genome reveals the origin and history of an ancient cell lineage. Science. 2014; 343:437–40.

28. Kwon YM, Gori K, Park N, Potts N, Swift K, Wang J, et al. Evolution and lineage dynamics of a transmissible cancer in Tasmanian devils. PLoS Biol. 2020; 18:e3000926.

29. Miller W, Hayes VM, Ratan A, Petersen DC, Wittekindt NE, Miller J, et al. Genetic diversity and population structure of the endangered marsupial Sarcophilus harrisii (Tasmanian devil). Proc Natl Acad Sci U S A. 2011; 108:12348–53.

30. Murchison EP, Schulz-Trieglaff OB, Ning Z, Alexandrov LB, Bauer MJ, Fu B, et al. Genome sequencing and analysis of the Tasmanian devil and its transmissible cancer. Cell. 2012; 148:780–91.

31. Deakin JE, Bender HS, Pearse AM, Rens W, O’Brien PC, Ferguson-Smith MA, et al. Genomic restructuring in the Tasmanian devil facial tumour: chromosome painting and gene mapping provide clues to evolution of a transmissible tumour. PLoS Genet. 2012; 8:e1002483.

32. Taylor RL, Zhang Y, Schoning JP, Deakin JE. Identification of candidate genes for devil facial tumour disease tumourigenesis. Sci Rep. 2017; 7:8761.

33. Skazina M, Odintsova N, Maiorova M, Ivanova A, Vainola R, Strelkov P. First description of a widespread Mytilus trossulus-derived bivalve transmissible cancer lineage in M. trossulus itself. Sci Rep. 2021; 11:5809.

34. Metzger MJ, Villalba A, Carballal MJ, Iglesias D, Sherry J, Reinisch C, et al. Widespread transmission of independent cancer lineages within multiple bivalve species. Nature. 2016; 534:705–9.

35. Yonemitsu MA, Giersch RM, Polo-Prieto M, Hammel M, Simon A, Cremonte F, et al. A single clonal lineage of transmissible cancer identified in two marine mussel species in South America and Europe. Elife. 2019; 8.

36. Tissot S, Meliani J, Boutry J, Brazier L, Tokolyi J, Roche B, et al. De novo evolution of transmissible tumours in hydra. Proc Biol Sci. 2024; 291:20241636.

37. Boutry J, Buysse M, Tissot S, Cazevielle C, Hamede R, Dujon AM, et al. Spontaneously occurring tumors in different wild-derived strains of hydra. Scientific Reports. 2023; 13.

38. Deycmar S, Gomes B, Charo J, Ceppi M, Cline JM. Spontaneous, naturally occurring cancers in non-human primates as a translational model for cancer immunotherapy. J Immunother Cancer. 2023; 11.

39. Dewi FN, Cline JM. Nonhuman primate model in mammary gland biology and neoplasia research. Laboratory Animal Research. 2021; 37.

40. Begley DA, Krupke DM, Sundberg JP, Jocoy EL, Richardson JE, Neuhauser SB, et al. The Mouse Models of Human Cancer database (MMHCdb). Disease Models & Mechanisms. 2023; 16.

41. Johnson LE, Becker JT, Dubovsky JA, Olson BM, McNeel DG. Prostate carcinoma in transgenic Lewis rats - a tumor model for evaluation of immunological treatments. Chin Clin Oncol. 2013; 2.

42. Miller JL, Bartlett AP, Harman RM, Majhi PD, Jerry DJ, Van de Walle GR. Induced mammary cancer in rat models: pathogenesis, genetics, and relevance to female breast cancer. J Mammary Gland Biol Neoplasia. 2022; 27:185–210.

43. Kasumi E, Chiba M, Kuzumaki Y, Kuzuoka H, Sato N, Takahashi B. Development and Characterization of a Cancer Cachexia Rat Model Transplanted with Cells of the Rat Lung Adenocarcinoma Cell Line Sato Lung Cancer (SLC). Biomedicines. 2023; 11:2824.

44. Duderstadt EL, McQuaide SA, Sanders MA, Samuelson DJ. Chemical carcinogen-induced rat mammary carcinogenesis is a potential model of p21-activated kinase positive female breast cancer. Physiol Genomics. 2021; 53:61–8.

45. Jia Y, Wang Y, Dunmall LSC, Lemoine NR, Wang P, Wang Y. Syrian hamster as an ideal animal model for evaluation of cancer immunotherapy. Frontiers in Immunology. 2023; 14.

46. Wang Z, Cormier RT. Golden Syrian Hamster Models for Cancer Research. Cells. 2022; 11.

47. Aizawa K, Liu C, Veeramachaneni S, Hu K-Q, Smith DE, Wang X-D. Development of ferret as a human lung cancer model by injecting 4-(N-methyl-N-nitrosamino)-1-(3-pyridyl)-1-butanone (NNK). Lung Cancer. 2013; 82:390–6.

48. Sprague J. The Zebrafish Information Network (ZFIN): a resource for genetic, genomic and developmental research. Nucleic Acids Research. 2001; 29:87–90.

49. Merlo DF, Rossi L, Pellegrino C, Ceppi M, Cardellino U, Capurro C, et al. Cancer incidence in pet dogs: findings of the Animal Tumor Registry of Genoa, Italy. J Vet Intern Med. 2008; 22:976–84.

50. Rafalko JM, Kruglyak KM, McCleary-Wheeler AL, Goyal V, Phelps-Dunn A, Wong LK, et al. Age at cancer diagnosis by breed, weight, sex, and cancer type in a cohort of more than 3,000 dogs: Determining the optimal age to initiate cancer screening in canine patients. PLOS ONE. 2023; 18:e0280795.

51. R. Graf KGMHKWASFMWDMFGGFVOAP. Swiss Feline Cancer Registry: A Retrospective Study of the Occurrence of Tumours in Cats in Switzerland from 1965 to 2008. Journal of Comparative Pathology. 2015; 153:266–77.

52. Kanfer S, Reavill DR. Cutaneous neoplasia in ferrets, rabbits, and guinea pigs. Vet Clin North Am Exot Anim Pract. 2013; 16:579–98.

53. Vasconcelos J, Pires MDA, Alves A, Vieira-Pinto M, Saraiva C, Cardoso L. Neoplasms in Domestic Ruminants and Swine: A Systematic Literature Review. Veterinary Sciences. 2023; 10:163.

54. Bertzbach L, Conradie A, You Y, Kaufer B. Latest Insights into Marek’s Disease Virus Pathogenesis and Tumorigenesis. Cancers. 2020; 12:647.

55. Nasir L, Campo MS. Bovine papillomaviruses: their role in the aetiology of cutaneous tumours of bovids and equids. Vet Dermatol. 2008; 19:243–54.

56. Duke EG, Harrison SH, Moresco A, Trout T, Troan BV, Garner MM, et al. A Multi-Institutional Collaboration to Understand Neoplasia, Treatment and Survival of Snakes. Animals (Basel). 2022; 12.

57. De Guise S, Lagace A, Beland P. Tumors in St. Lawrence beluga whales (Delphinapterus leucas). Vet Pathol. 1994; 31:444–9.

58. Martineau D, Lemberger K, Dallaire A, Labelle P, Lipscomb TP, Michel P, et al. Cancer in wildlife, a case study: beluga from the St. Lawrence estuary, Quebec, Canada. Environ Health Perspect. 2002; 110:285–92.

59. Fox KA, Wootton SK, Quackenbush SL, Wolfe LL, Levan IK, Miller MW, et al. Paranasal sinus masses of Rocky Mountain bighorn sheep (Ovis canadensis canadensis). Vet Pathol. 2011; 48:706–12.

60. Integrated Canine Data Commons. In: Institute NC, editor.: National Institute of Health.

61. Rochman ND, Wolf YI, Koonin EV. Deep phylogeny of cancer drivers and compensatory mutations. Commun Biol. 2020; 3:551.

62. Burningham Z, Hashibe M, Spector L, Schiffman JD. The Epidemiology of Sarcoma. Clinical Sarcoma Research. 2012; 2:14.

63. Clauss M, Müller DWH. Putting zoo animal cancer into perspective. Zoo Biology. 2024; 43:15–21.

64. Pawłowski B, Żelaźniewicz A. The evolution of perennially enlarged breasts in women: a critical review and a novel hypothesis. Biol Rev Camb Philos Soc. 2021;96(6):2600–2615. doi:10.1111/brv.12778

65. Walser T, Cui X, Yanagawa J, Lee JM, Heinrich E, Lee G, et al. Smoking and lung cancer: the role of inflammation. Proc Am Thorac Soc. 2008;5(8):811–815. doi:10.1513/pats.200809-100TH

66. Lopez-Garcia C, Klein AM, Simons BD, Winton DJ. Intestinal stem cell replacement follows a pattern of neutral drift. Science. 2010;330(6005):822–825. doi:10.1126/science.1196236

67. Aupperle-Lellbach H, Grassinger JM, Floren A, Törner K, Beitzinger C, Loesenbeck G, et al. Tumour Incidence in Dogs in Germany: a Retrospective Analysis of 109,616 Histopathological Diagnoses (2014–2019). Journal of Comparative Pathology. 2022; 198:33–55.

68. Gustafson DL, Duval DL, Regan DP, Thamm DH. Canine sarcomas as a surrogate for the human disease. Pharmacology & Therapeutics. 2018; 188:80–96.

69. Huang Z, Whelan CV, Dechmann D, Teeling EC. Genetic variation between long-lived versus short-lived bats illuminates the molecular signatures of longevity. Aging (Albany NY). 2020;12(16):15962–15977. doi:10.18632/aging.103725

70. Wirthlin M, Lima NCB, Guedes RLM, Vasconcelos ATR, Prosdocimi F, Mello CV, et al. Parrot genomes and the evolution of heightened longevity and cognition. Curr Biol. 2018;28(24):4001–4008.e7. doi:10.1016/j.cub.2018.10.050

71. Tollis M, Schneider-Utaka AK, Maley CC. The Evolution of Human Cancer Gene Duplications across Mammals. Mol Biol Evol. 2020; 37:2875–86.

72. Bartas M, Brazda V, Volna A, Cerven J, Pecinka P, Zawacka-Pankau JE. The Changes in the p53 Protein across the Animal Kingdom Point to Its Involvement in Longevity. Int J Mol Sci. 2021; 22.

73. Foley NM, Hughes GM, Huang Z, Clarke M, Jebb D, Whelan CV, et al. Growing old, yet staying young: The role of telomeres in bats’ exceptional longevity. Science Advances. 2018; 4:eaao0926.

74. Iranzo J, Gruenhagen G, Calle-Espinosa J, Koonin EV. Pervasive conditional selection of driver mutations and modular epistasis networks in cancer. Cell Reports. 2022; 40:111272.

75. Rhie A, McCarthy SA, Fedrigo O, Damas J, Formenti G, Koren S, et al. Towards complete and error-free genome assemblies of all vertebrate species. Nature. 2021; 592:737–46.

76. ResearchRabbit.ai. R. Internet; Available from: https://www.researchrabbit.ai/.

77. Mattiuzzi C, Lippi G. Current Cancer Epidemiology. Journal of Epidemiology and Global Health. 2019; 9:217.

78. Shahriari M, Namavari M, Ziyaeyan M, Etemadfar P. Relationship between Human Herpes Virus Type 6 and Childhood T-Cell Leukemia-Lymphoma. Iranian Red Crescent Medical Journal (IRCMJ). 2022; 24:-.

79. Comprehensive and Integrated Genomic Characterization of Adult Soft Tissue Sarcomas. Cell. 2017; 171:950-65.e28.

80. Ahmadipour M, Kitzwögerer M, Trautinger F. Retrospective study of postoperative survival of keratinocyte-derived skin cancer patients at the end of life. J Dtsch Dermatol Ges. 2024; 22:1213–8.

81. Bekampytė J, Savukaitytė A, Bartnykaitė A, Ugenskienė R, Žilienė E, Inčiūra A, et al. TIRAP Rs8177376, Rs611953, Rs3802814, and Rs8177374 Polymorphisms and Their Association with Cervical Cancer Phenotype and Prognosis. Genes (Basel). 2022; 13.

82. Cheishvili D, Stefanska B, Yi C, Li CC, Yu P, Arakelian A, et al. A common promoter hypomethylation signature in invasive breast, liver and prostate cancer cell lines reveals novel targets involved in cancer invasiveness. Oncotarget. 2015; 6:33253–68.

83. Khelf S, Guedjali L, Souad H, Dalila S. Molecular Classification of Breast Cancer in the Region of Constantine: An Epidemiological Study. Journal of Nanosciences Research & Reports. 2020:1–3.

84. Ma Z, Zou X, Yan Z, Chen C, Chen Y, Fu A. Preliminary Analysis of Cervical Cancer Immunotherapy. Am J Clin Oncol. 2022; 45:486–90.

85. Markiewicz A, Sigorski D, Markiewicz M, Placek WJ, Owczarczyk-Saczonek AB. mRNA expression of caspase 14 in skin epithelial malignancies. Postepy Dermatol Alergol. 2023; 40:315–20.

86. Baba AI CC. Comparative Oncology. Bucharest (RO): The Publishing House of the Romanian Academy; 2007.

87. Hosseini H, Heydari S, Hushmandi K, Daneshi S, Raesi R. Bone tumors: a systematic review of prevalence, risk determinants, and survival patterns. BMC Cancer. 2025; 25:321.

88. Price M, Neff C, Nagarajan N, Kruchko C, Waite KA, Cioffi G, et al. CBTRUS Statistical Report: American Brain Tumor Association & NCI Neuro-Oncology Branch Adolescent and Young Adult Primary Brain and Other Central Nervous System Tumors Diagnosed in the United States in 2016–2020. Neuro-Oncology. 2024; 26:iii1–iii53.

89. Howlader N, Noone AM, Krapcho M, Miller D, Brest A, Yu M, Ruhl J, Tatalovich Z, Mariotto A, Lewis DR, Chen HS, Feuer EJ, Cronin KA, editors. SEER Cancer Statistics Review, 1975-2018. Bethesda, MD: National Cancer Institute; 2021 [cited 2025 Oct 10]. Available from: https://seer.cancer.gov/csr/1975_2018/

90. Sung H, Ferlay J, Siegel RL, Laversanne M, Soerjomataram I, Jemal A, Bray F. Global Cancer Statistics 2020: GLOBOCAN Estimates of Incidence and Mortality Worldwide for 36 Cancers in 185 Countries. CA Cancer J Clin. 2021;71(3):209–249. doi:10.3322/caac.21660. Available from: https://acsjournals.onlinelibrary.wiley.com/doi/10.3322/caac.21660

