## Supplementary Figures for "Harmonized nonhuman cancer database"

### Cancer origins across vertebrates

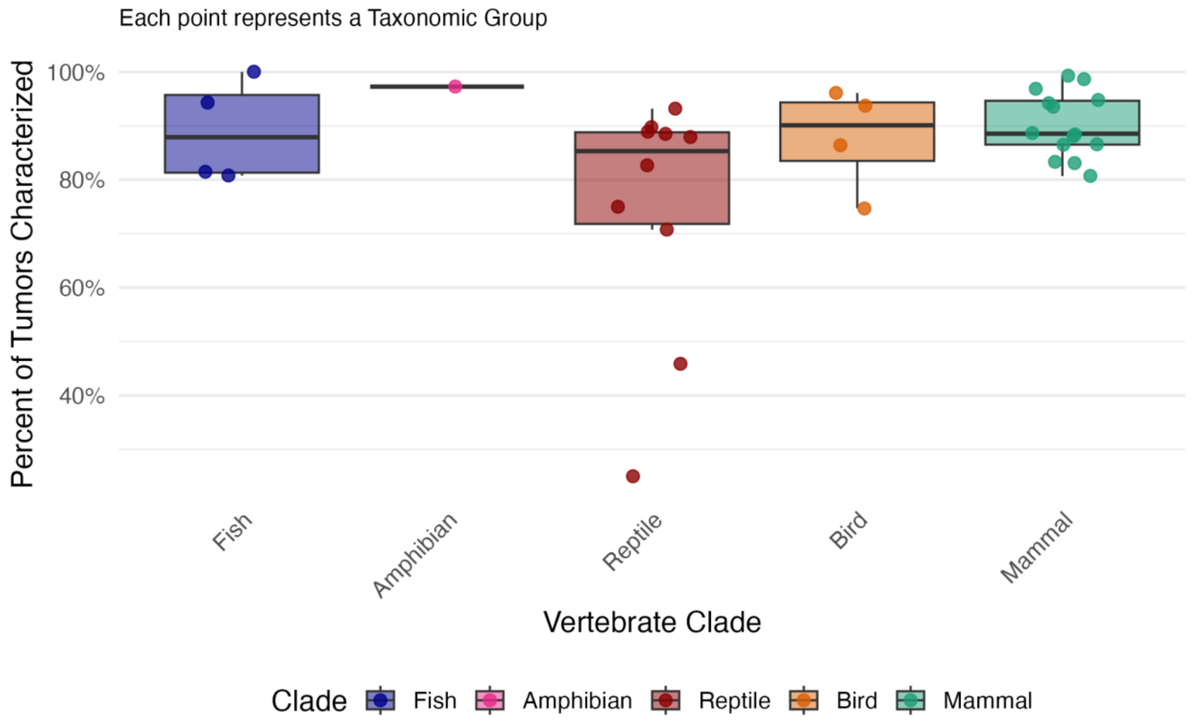


**Supplementary Figure 1.** Tumor characterization is similarly supported across major taxonomic groups.

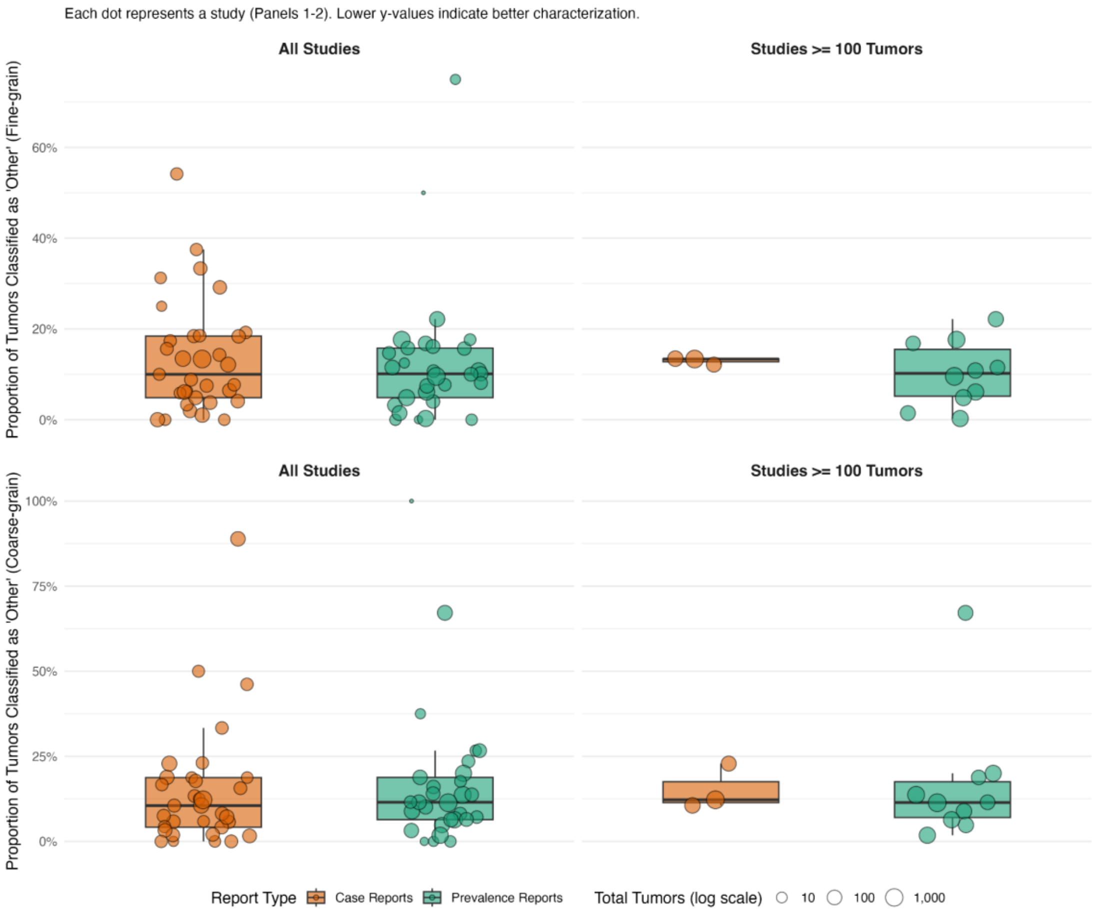


**Supplementary figure 2.** Top row reflects 16-category body-site classification, Fine-grain (see main text). Bottom row reflects 3-category classification into carcinoma, sarcoma or lymphohematopoietic cancer, Coarse-grain. When metadata is absent, the tumor is labelled, “Other”. Bubble size corresponds to the log number of tumors in each study or taxonomic group, respectively.
